# From haptic exploration to object perception in 3D

**DOI:** 10.64898/2026.10.12.758479

**Authors:** Anna Metzger, Matteo Toscani

## Abstract

Active touch determines which parts of an object are sampled and therefore which sensory information becomes available for perception. However, little is known about how the three-dimensional geometry of an object guides exploratory hand movements and how the information acquired during exploration contributes to perceptual decisions. Here, we investigated these questions in two experiments in which blindfolded participants explored 3D-printed replicas of bell-peppers while their hand movements were recorded using motion tracking. The task of participants’ was to identify the reference object among two comparison objects (target & competitor). In the first experiment, objects were presented in a fixed orientation, whereas in the second, object orientation varied across presentations to disrupt the correspondence between hand configuration and object geometry. We first trained a neural network to predict the spatial distribution of touch directly from three-dimensional surface structure. The model successfully predicted touch locations on previously unseen objects, outperforming a generic spatial exploration prior in both experiments. Geometric analyses revealed that haptic saliency was associated with specific combinations of local surface curvatures rather than simply with the overall magnitude of curvature. We then examined whether the information acquired during exploration could predict subsequent recognition choices, considering global and local surface geometry and potential memory decay of acquired object representations. Models integrating global and local information across the complete exploration reliably identified the objectively correct object and correct participants’ choices. In contrast, restricting the comparison to temporally proximal portions of successive explorations reduced objective recognition accuracy but increased correspondence with participants’ errors. Overall, our findings suggest that haptic exploration is at least to some extent driven by object-specific geometry, but that the information acquired is not necessarily integrated optimally into perceptual decisions. Instead, memory limitations and potentially incomplete integration may constrain how sampled three-dimensional shape information contributes to recognition.

## Introduction

Perception by touch is inseparable from action. Active touch differs fundamentally from passive cutaneous stimulation because observers actively move the hand to acquire information about object properties (1). During haptic object recognition, hand movements are not arbitrary: distinct exploratory procedures are selected depending on the information being sought, and these movements are necessary and optimal for extracting specific object properties (2). The hand therefore continuously determines where an object is contacted, which fingers are involved and how contact unfolds over time. These exploratory actions constrain the sensory evidence available to the nervous system and can consequently shape the representation acquired of an object’s three-dimensional structure. A central question for active touch is therefore not only how local tactile signals encode shape, but also whether and why some surface regions are sampled more than others.

Vision provides a useful conceptual analogy. Computational accounts of visual saliency propose that local image features can be integrated into a spatial priority or saliency map, such that some locations are preferentially selected for further processing (3). Visual selection is not determined by stimulus salience alone; where observers look also depends on behavioural goals and the demands of the current task (4–7). This distinction motivates an analogous question in touch: whether the geometry of a three-dimensional object gives rise to a spatial distribution of haptic saliency, such that some surface regions are more likely than others to be selected during exploration. To our knowledge, we provided the first direct evidence for such stimulus-driven haptic saliency in previous work, showing that the spatial distribution of touch on textured relief surfaces could be predicted from local surface properties (8,9). Those studies, however, aimed to investigate information selection in non-uniform textures, and hence concerned surfaces that were effectively represented over an approximately planar substrate. Fully three-dimensional objects introduce additional geometric structure that is absent, or only weakly expressed, in such 2D surface representations. In particular, local shape can be described in terms of principal curvatures and derived quantities such as mean curvature, Gaussian curvature, shape index and curvedness, which distinguish convex, concave and saddle-like regions. Here, we ask whether haptic saliency extends to the exploration of fully three-dimensional objects, for which potentially informative surface regions are distributed over the entire object and can be characterized by this richer local geometry. A separate question is whether the sensory evidence acquired through this exploration is actually related to the observer’s perceptual decision. In vision, active sampling and perceptual choice are tightly linked: eye movements determine which parts of the scene are sampled at high resolution, and models of visual search treat successive fixations as a means of gathering sensory evidence for a later perceptual decision (10,11). A similar logic applies to touch. Manual exploratory procedures are adapted to the object property being judged, and constraining exploration impairs haptic identification, indicating that the pattern of sampling determines which sensory information becomes available for recognition (2,12,13). Furthermore, our preliminary findings show that for non-uniform object surfaces there are foveation-like aspects in haptic exploration, similar to vision, indicating active selective sampling with the higher resolving finger (14). Yet it remains less clear how the sampled sensory input is related to perceptual choices.

Haptic exploration of 3D objects provides both cutaneous information, arising from the activation of afferents in the skin, and proprioceptive information about the spatial configuration of the fingers and hand (15,16). Previous behavioural studies have shown that these sources of information jointly contribute to shape discrimination and object recognition (17). This is supported by physiological findings that neurons in the somatosensory cortex respond to both cutaneous and proprioceptive stimulation, with some exhibiting interactions between these signals, such that responses to cutaneous stimulation are modulated by hand conformation and vice versa (18). In addition, haptic shape judgments are sensitive to local surface geometry, including curvature-based descriptions of shape (19–21). Despite the fact that haptic exploration could in principle recover the entire 3D structure of the object, haptic object recognition is influenced by object orientation (22). This raises the question to what extent perceptual choices are related to the sampled information independent of the posture of the exploring hand.

In the present study, we thus asked two related questions. First, is haptic exploration of fully three-dimensional objects predictable from object geometry, and which local surface properties are associated with locations that are preferentially touched? Second, does global and local information acquired during exploration predict subsequent object discrimination? We modelled global information as positions contacted on the object surface, and local information as geometric properties available at touched locations on the object. We additionally asked how the temporal availability of this information influenced recognition predictions. A decision need not reflect an optimal integration of every contact acquired during exploration: information sampled at different stages may receive different weights. Indeed, Metzger and colleagues showed that, when comparing two objects, repeated estimates of the first object were weighted approximately equally, whereas estimates of the second object obtained shortly after the exploration of the first object contributed more strongly to the perceptual decision than later estimates (23–25). We addressed these questions in two experiments in which blindfolded participants explored 3D-printed bell-peppers (1 reference and 2 comparisons) while recording hand movements with motion tracking. To test whether the sampled information predicts perceptual choices independent of hand posture we conducted two experiments. In Experiment 1, reference and comparison objects were presented in the same orientation; in Experiment 2, comparison objects were randomly rotated around the vertical axis relative to the reference, disrupting the correspondence between hand configuration and object geometry. We first modelled where participants touched and characterised the geometric structure of the resulting haptic-saliency maps. We then tested whether recognition choices could be related to object-centred contact position or local surface shape. We also compared models using the complete exploration with models retaining different portions of the initial target and subsequent comparison explorations, including conditions in which a complete representation of the target was compared with early information from the later objects and more temporally local comparisons between the end of target exploration and the beginning of comparison exploration. In this way, we sought to characterise the full pipeline from active sampling of three-dimensional shape, through the sensory information acquired during exploration, to the temporal use of that information in the final recognition decision.

## Material and methods

### Participants and ethics

Nine participants, all righthanded volunteered to take part in Experiment 1 (7 female, 2 male; mean age = 27.2 years, SD = 7.4, range = 21–41 years) and ten participants in Experiment 2 (8 female, 2 male, mean age = 27.8 years, SD = 5.2, range = 21–35 years). All participants provided written informed consent before taking part. The experiments were conducted in accordance with the Declaration of Helsinki and were approved by the Ethics Committee of Bournemouth University (ID: 59969). Participants were compensated for their participation with course credits or vouchers (1 credit / £10 voucher per hour).

### Stimuli

The stimuli were ten differently shaped bell-peppers (Capsicum annuum) as previously used by Norman and colleagues (26). Bell peppers have been used extensively as natural-object stimuli in studies of visual and haptic shape perception because individual exemplars are novel to participants, they share the same broad object category and have roughly the same volume while differing substantially in their detailed three-dimensional surface geometry (26–28). Hence, recognition cannot be based simply on category membership or simple feature like size, but requires sensitivity to the spatial configuration of surface structure and local surface geometry. The bell-pepper shapes are also especially well suited to the present study because they are sufficiently large to prompt hand movements for exploration (as opposed to just enclose them in the hand as could be expected for smaller objects) and their surfaces contain a heterogeneous mixture of locally convex, concave and saddle-like regions. Norman et al. (28) have argued that this spatial heterogeneity of local curvature is important for the recognition of natural three-dimensional objects. For the present experiments, we used digital models of the peppers (provided with courtesy by J. Farley Norman) to 3D-print real size replicas using cornstarchbased polylactic acid and a MakerBot Replicator (5^th^ Gen) Desktop 3D Printer. Their weight was between 60 and 100 g. As the haptic stimuli were fixed to a pedestal for exploration and were never lifted, variations in the weight were non-informative for the perceptual task. In Experiment 1, the objects were presented in a fixed orientation, so repeated presentations of the same object preserved the relationship between object orientation and the external coordinate frame. In Experiment 2, all objects were presented at different, predefined, randomly assigned orientations around the vertical axis. Four orientations were possible, corresponding to rotations of 0°, 90°, 180° and 270° (0° corresponds to the front of the pepper facing the participant), allowing the relationship between the explored object surface and the observer’s hand to vary across presentations.

### Apparatus and motion tracking

Hand movements were recorded using a Qualisys motion-capture system (Qualisys AB, Gothenburg, Sweden) comprising nine cameras, operating at a sampling frequency of 100 Hz. Prior to each data collection, the motion tracking system was calibrated using a calibration wand and an L-shaped reference frame to establish the spatial coordinate system and calibrate the camera arrangement (mean residual error **>** 0.25 mm). Seventeen reflective markers were attached to the participant’s right hand, enabling automatic tracking. For analysis only three-dimensional positions of the markers on the fingertips were used. The physical position of each pepper (in each orientation for Experiment 2) was registered prior to the experiment by attaching markers to the **object**. To establish the spatial correspondence between the physical objects and their digital 3D models, **t**he recorded marker positions and the corresponding 3D object mesh were manually rotated to align in Blender (Blender Foundation, Amsterdam, the Netherlands), thereby registering the digital object geometry to the motion-capture coordinate system **(29)**. In Experiment 2, this procedure was repeated for each object orientation. The resulting alignment allowed the recorded fingertip trajectories to be related to the corresponding locations on the object’s 3D surface.

### Procedure

On each trial, participants explored three successively presented objects with the right hand. The first object served as the reference. It was followed by two comparison objects, one of which was the same pepper as the reference (target) and the other a different pepper (competitor). Participants were required to indicate which of the two comparison objects matched the reference.

Participants were allowed to explore each object for a constrained period of either 2 or 5 s. Exploration duration varied across trials, such that participants did not know in advance how long a particular exploration would last. The same duration applied to the successive object presentations within a trial. A tone signalled the end of the exploration time and prompted participants to remove their hand from the. To prevent visual access to the objects during exploration, participants wore disposable surgical face masks positioned over their eyes as blindfolds. The masks were secured behind the ears using their elastic ear loops, as in normal use, and the integrated adjustable nose clip was fitted around the nose to prevent participants from seeing through gaps beneath the mask. Disposable masks were used for hygiene purposes. by a blindfold. We used facemasks as blind folds as they. The order of target and competitor comparisons was randomly varied across trials.

### Estimating contact

For each time sample, the Euclidean distance between each fingertip marker and all vertices of the corresponding pepper surface, transformed into motion-tracking coordinates, was computed. A fingertip was classified as being in contact with the object when its distance to the nearest surface vertex was ≤ 2 cm. For samples classified as contact, the nearest surface vertex was taken as the corresponding contact location. These locations were retained as the spatial record of exploration and, where required, associated with the identity of the contacting finger.

### Neural-network prediction of touch

To test whether the spatial pattern of haptic exploration could be predicted from object geometry, we represented each pepper directly as a sampled three-dimensional surface and trained a local point-cloud network to assign a saliency value to each surface location. Predictions were defined only at locations on the object surface. The architecture (Figure 1G) was inspired by the shared point-wise transformations and symmetric pooling used in PointNet, together with the use of local metric neighbourhoods in PointNet++ (30,31). Importantly, the network was not given curvature or other hand-engineered shape descriptors; these were calculated only in subsequent analyses used to characterise the geometry associated with the learned predictions. Each pepper retained its native triangular mesh representation. To obtain a fixed-resolution representation suitable for the network and for comparison of touch distributions, we independently sampled 8192 locations from each mesh surface. Triangles were selected in proportion to their surface area, and points were sampled within the selected triangles using barycentric coordinates, yielding an approximately uniform surface-area sampling. The resulting 8192 locations were analysis samples rather than native mesh vertices and were not assumed to correspond across different peppers. The number of sampled surface locations was fixed across objects so that the network operated on representations of identical dimensionality, irrespective of differences in the number or tessellation of vertices in the original reconstructed meshes. These 8192 points defined the locations at which touch saliency was represented and predicted.

**Figure 1.**
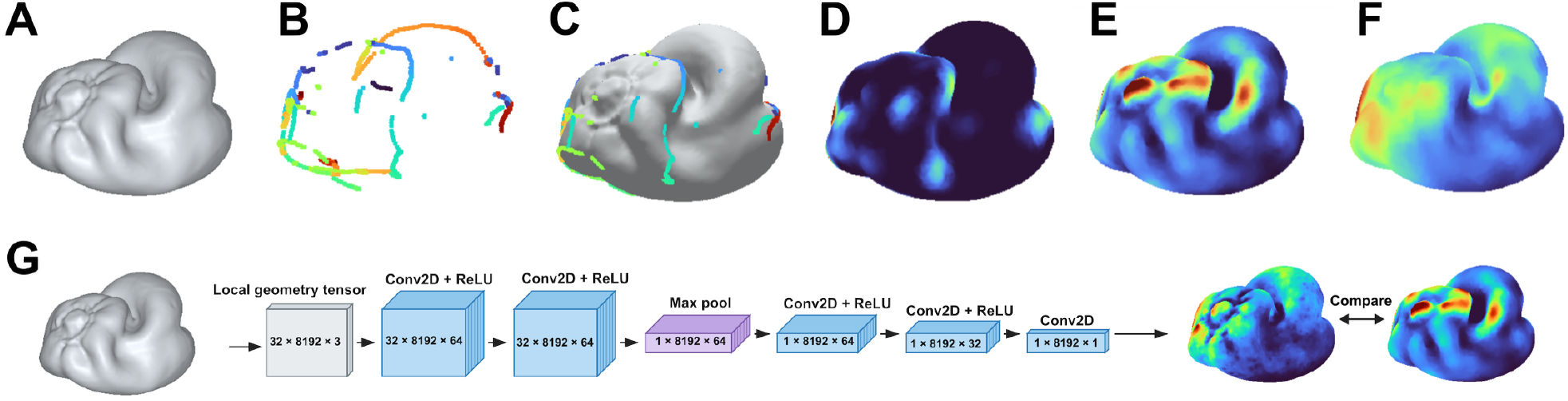
Surface representation of haptic exploration and DNN model. **(A)** Example three-dimensional pepper surface. **(B)** Example fingertip contact trajectories during exploration, shown without the object surface. Colours indicate progression along the recorded trajectories. **(C)** The same contact trajectories shown on the corresponding pepper surface. **(D)** Trial-level touch-density map obtained by projecting the recorded contacts onto the surface and spreading each contact over nearby surface locations with distance-dependent weighting. **(E)** Mean touch-density map for that pepper, obtained by averaging the trial-level touch-density maps. **(F)** Generic spatial prior, estimated from touch distributions on the other peppers and spatially interpolated onto the target pepper; this captures spatial regularities in exploration that are shared across objects. Warmer colours in D–F indicate greater touch density or probability. **(G)** Architecture of the geometry-based network. Each pepper was represented by 8192 sampled surface locations, with the local geometry around each location described by the relative XYZ coordinates of its 32 nearest surface neighbours, producing a 32 x 8192 x 3 input tensor. Two shared 1×1 convolutional layers with 64 channels and ReLU activations extracted features independently from each neighbour. Max pooling across the 32 neighbours produced one 64-dimensional local-geometry descriptor for each surface location. A point-wise prediction head then transformed these descriptors through 64- and 32-channel layers to a single geometry-dependent value at each of the 8192 surface locations. These values were combined with the generic spatial prior to generate the predicted haptic-saliency map, which was compared with the observed touch-density map.

In Experiment 1, all peppers were analysed in their fixed presentation orientation. In Experiment 2, contacts were first detected on the mesh in the orientation actually presented on that trial, then mapped onto a common canonical orientation for analysis. This preserved orientation-specific contact detection while placing all predictions in a common object-centred reference frame.

Touch was represented on the same 8192-point surface. Finite fingertip samples were retained at the original 100-Hz sampling rate, so repeated samples at the same location contributed to the representation of exploration. Samples lying within a 2-cm surface-contact criterion, scaled relative to a 15-cm reference object diameter, were projected to the nearest compatible mesh triangle. For each presentation, the projected contact locations were converted into a continuous touch-density map by spreading the contribution of each contact over nearby surface points with a Gaussian weighting function. Points closest to a contact received the greatest weight, with the contribution decreasing smoothly with distance (sigma = 5 mm). The contributions from all contacts were then summed and the resulting map normalised to sum to one. A separate binary map marked the sampled surface point nearest to at least one observed contact. For model training, presentation-level density maps were first normalised separately and then averaged within each participant-object combination, so that longer presentations could not dominate solely because they contained more recorded frames. For each participant– pepper combination, the continuous touch-density maps from the relevant presentations were averaged to form the primary training target. In parallel, we retained a binary contact mask indicating whether each sampled surface location had been contacted at least once across those presentations. This binary mask was used only for the auxiliary ranking loss and for AUC evaluation, not as the main prediction target.

The geometric input to the network described the local surface shape around each of the 8192 sampled locations. For every sampled centre point **x**_*i*_, we identified its 32 nearest other sampled surface points in Euclidean three-dimensional distance. Each neighbour was expressed relative to its centre point,

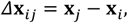

where Δ**x**_*ij*_ contains the three coordinate differences (*Δx, Δy, Δz*). Consequently, every one of the 8192 centre points was represented by a local 32 × 3 set of relative coordinates. Stacking all local neighbourhoods produced a 32 × 8192 × 3 input tensor. Expressing neighbours relative to the centre removes translation of the local patch and allows local surface configurations to be learned directly from raw coordinates. No absolute XYZ position, surface normal, principal curvature, mean curvature, Gaussian curvature, shape index or curvedness was supplied to the retained predictive model.

First, every three-dimensional neighbour offset was independently passed through a shared 1 × 1 two-dimensional convolution with 64 output channels followed by a rectified-linear activation (ReLU). Because the kernel was 1 × 1, the same set of learned weights was applied independently to the coordinate vector of every neighbouring point. A second shared 1 × 1 convolution with 64 output channels and ReLU was then applied. The representation at this stage therefore had dimensions 32 × 8192 × 64: for each of the 8192 surface locations, each of its 32 neighbours had been transformed into a learned 64-dimensional feature vector. The 32 neighbour representations were then collapsed using a maximum over the neighbour dimension, independently for each of the 64 learned feature channels. This changed the representation from 32 × 8192 × 64 to 1 × 8192 × 64. Max pooling is symmetric with respect to neighbour order, so the resulting 64-dimensional descriptor of a local surface patch does not depend on the arbitrary order in which its 32 neighbouring points are listed, following the permutation-invariant aggregation principle introduced in PointNet (30). The operation compresses the local neighbourhood but does not remove any of the 8192 surface locations. A point-wise prediction head was subsequently applied independently to each of the 8192 local descriptors. It comprised a 1 × 1 convolution mapping 64 channels to 64 channels followed by ReLU, a second 1 × 1 convolution mapping 64 channels to 32 channels followed by ReLU, and a final linear 1 × 1 convolution mapping 32 channels to a single scalar. The dimensional sequence of the complete network was therefore 32 × 8192 × 3 → 32 × 8192 × 64 → 32 × 8192 × 64 → 1 × 8192 × 64 → 1 × 8192 × 64 → 1 × 8192 × 32 → 1 × 8192 × 1.

This architecture was chosen because the reconstructed peppers are naturally represented as irregular surface point sets rather than dense Cartesian volumes. PointNet demonstrated that shared point-wise transformations followed by a symmetric aggregation function can operate directly on unordered point clouds without voxelisation (30). The present model additionally defines a local metric neighbourhood around every candidate surface location, reflecting the motivation of PointNet++ that local spatial relationships are needed to capture fine-scale geometric structure (31). Here, however, the architecture was deliberately kept shallow: its purpose was not object classification, but learning a local geometric descriptor at each surface point from which haptic saliency could be predicted.

To distinguish object-specific geometric information from generic spatial regularities in exploration, the network was trained as a residual correction to a fixed pepper-aspecific spatial prior. For each outer leave-one-pepper-out fold, the prior for a target pepper was constructed only from the other available peppers. Participant touch-density maps were first averaged within each source pepper so that every source object contributed equally. The 8192 canonical surface coordinates and their mean touch values were then pooled across source peppers. To construct the generic spatial prior for a target pepper, participant touch-density maps were first averaged within each of the other peppers, giving each source pepper equal weight. Because the 8192 sampled surface locations did not correspond point-for-point across different peppers, these mean maps could not be averaged directly by surface-point index. Instead, for each surface location on the target pepper, we identified the 200 nearest sampled locations from the other peppers in canonical XYZ space and calculated a distance-weighted mean of their touch values. Nearby source locations contributed more strongly than distant ones according to a Gaussian weighting function (sigma=0.05 in diameter-normalised coordinates). The resulting values were then normalised to sum to one over the target surface, yielding the generic spatial prior. During training, the prior for each training pepper was itself constructed from the other training peppers, excluding both that pepper and the outer held-out test pepper. For the held-out pepper, the prior was constructed from all non-held-out peppers. Touch observations from an object therefore never contributed to the prior used to predict that same object.

Let *r*_*i*_ denote the scalar output of the geometry network at surface location *i*, and let *M*_*i*_ denote the fixed generic spatial prior. Final unnormalised log saliency was defined as

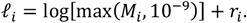

The final predicted surface distribution was obtained by a softmax over all 8192 locations. This residual formulation has a useful interpretation: if the geometry network outputs *r*_*i*_ = 0 everywhere, the final prediction is exactly the generic spatial prior. The network is therefore required only to learn deviations from generic spatial sampling that can be supported by the local geometry of a particular object.

Networks were trained for 60 epochs using minibatches of four participant-object maps and the Adam optimiser with a learning rate of 5 × 10^−4^. MATLAB’s adamupdate routine was used with its default first- and second-moment decay factors (*β*_1_ = 0.9 and *β*_2_ = 0.999) and numerical constant *ϵ* = 10^−8^. Training used a compound objective combining the continuous touch-density distribution with the binary contacted/non-contacted map. The density term was cross-entropy between the observed touch-density distribution *T* and the final predicted distribution *P*,

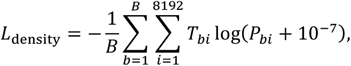

where *B* is the minibatch size. A second term encouraged contacted locations to receive higher final logits than non-contacted locations. For each example *b*, the mean final logit over contacted locations was denoted *µ*_+,*b*_ and the mean over non-contacted locations *µ*_−,*b*_. The ranking loss was a smooth margin loss,

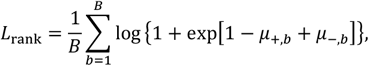

with a margin of 1. A small residual penalty discouraged unnecessarily large deviations from the generic prior,

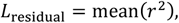

and an L2 penalty was applied to all learnable parameters *θ* of the two network components,

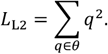

The complete training objective was

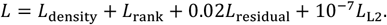

Thus, the density term trained the model to reproduce the continuous distribution of touch over the surface, the ranking term directly encouraged separation between contacted and non-contacted locations, and the residual penalty favoured the generic prior unless the learned geometry provided evidence for a systematic correction.

Generalisation was evaluated using leave-one-pepper-out cross-validation. In each fold, one pepper identity was removed completely from network training and from construction of all training priors. The trained network was then applied once to the geometry of the unseen pepper to generate its predicted saliency distribution. Predictive performance was quantified using the area under the receiver-operating-characteristic curve (AUC), calculated exactly from the ranks of predicted saliency values at contacted and non-contacted surface locations. Tied prediction values were assigned their mean rank. AUC was calculated separately for each participant who had explored the held-out object and then averaged within pepper. An AUC of 0.5 corresponds to chance ranking, whereas values above 0.5 indicate that locations receiving higher model saliency were more likely to have been contacted.

Two reference predictions were evaluated with the same AUC measure. The first was the pepper-aspecific generic spatial prior *M* described above. Because zero output from the geometry network reproduces this prior exactly, model performance above the generic prior provides a direct test of predictive information contributed by object-specific local geometry. The second reference quantified participant agreement. For each participant and held-out pepper, the predictor was the mean touch-density map of the *other* participants who had explored that same pepper. This same-object, leave-one-participant-out map estimates the reproducible object-specific structure present in human exploration and therefore serves as an empirical upper reference for the geometry model; performance approaching this level would indicate that the model captures most of the reproducible structure available in human touch patterns.

### Geometric analysis of predicted haptic saliency

To determine which aspects of local three-dimensional surface geometry were associated with predicted haptic saliency, we related the neural network’s voxel-wise touch probabilities to a set of curvature-based shape descriptors (Figure 2) computed from the same voxelized object representation. Surface voxels were first identified as occupied voxels with at least one empty voxel in their 26-voxel neighbourhood. For each surface voxel, local geometry was estimated from the occupied voxels within a 3 × 3 × 3 neighbourhood. The neighbouring points were centred on their centroid, and principal-component analysis was used to estimate the local surface normal. The points were then expressed relative to the corresponding tangent plane, and a local quadratic surface was fitted:

**Figure 2.**
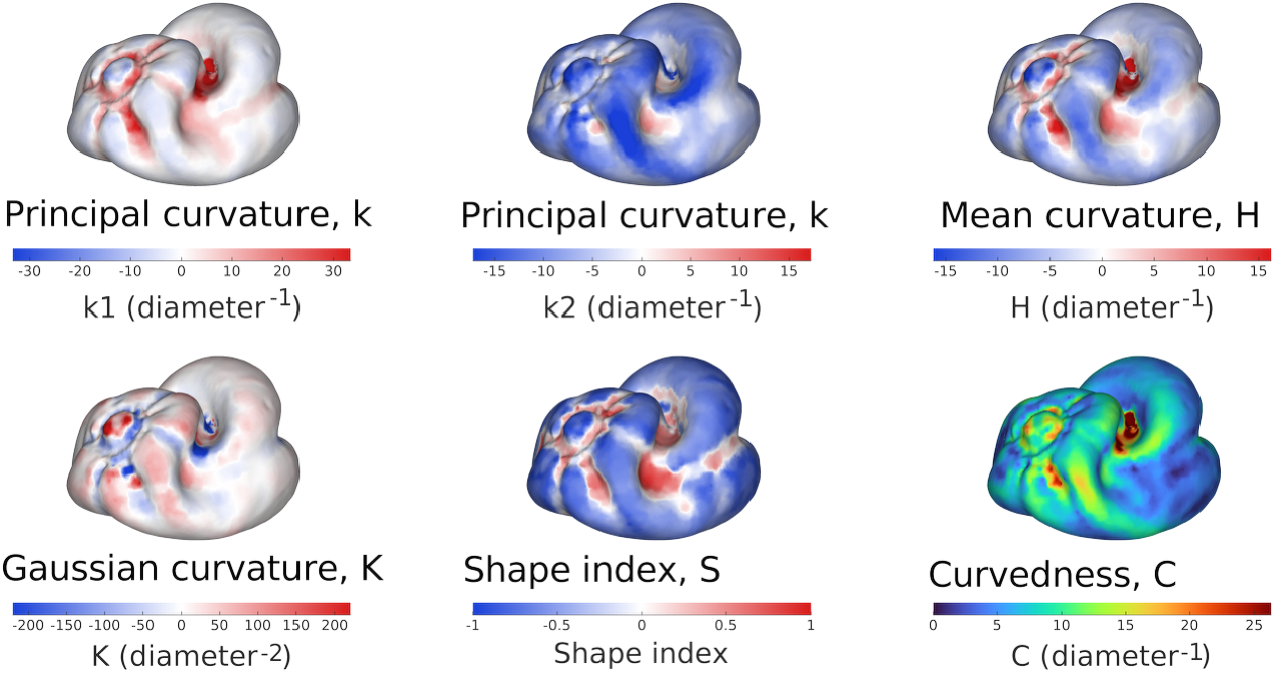
Local surface-geometry measures used to characterise haptic saliency. Example maps are shown for one pepper, with colour indicating the value of each geometric descriptor at each sampled surface location. The six measures were derived from the local principal curvatures: the maximum principal curvature k1, minimum principal curvature k2, mean curvature, Gaussian curvature, shape index, and curvedness C. Signed measures are shown with diverging colour scales centred on zero, whereas curvedness, which is non-negative, is shown with a sequential colour scale. Curvature values are expressed in diameter-normalised units. These descriptors were not provided to the neural network; they were computed separately to quantify which aspects of local surface geometry were associated with human touch patterns and with model-predicted haptic saliency.

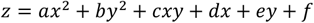

where x and y denote coordinates in the local tangent plane and z denotes displacement along the estimated surface normal. At least six points were required for the fit. The second-order terms of this fitted surface were used to form the local curvature matrix:

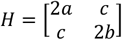

The two eigenvalues of this matrix were taken as estimates of the two principal curvatures, k_1_ and k_2_.

From these principal curvatures, four additional geometric descriptors were derived. Mean curvature was defined as:

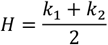

and describes the signed average bending of the surface.

Gaussian curvature was defined as:

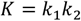

and characterizes the qualitative type of local surface geometry. Positive values indicate that the two principal curvatures have the same sign, as in locally dome- or bowl-like regions, whereas negative values indicate curvatures of opposite sign, corresponding to saddle-like regions. Values near zero indicate that curvature is small in at least one principal direction.

Shape index was calculated as:

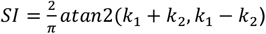

providing a continuous description of local shape based on the relative signs and magnitudes of the principal curvatures. Finally, curvedness was defined as:

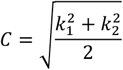

which describes the overall magnitude of local curvature irrespective of its sign. Thus, six local surface measures were considered in total: the two principal curvatures, mean curvature, Gaussian curvature, shape index and curvedness.

For each voxelized object, these six measures were stored as spatial feature maps and compared with the touch-probability map predicted by the trained neural network. For each descriptor separately, we calculated the Spearman rank correlation between the geometric value and the predicted probability of touch across voxels for which the descriptor was defined. The analysis therefore asked whether locations assigned higher predicted saliency by the network tended systematically to exhibit particular local geometric properties. A positive correlation indicates that predicted touch probability increased with the value of the corresponding descriptor, whereas a negative correlation indicates that higher predicted touch probability was associated with lower descriptor values. Correlations were calculated separately for each object and then summarized across objects to assess which geometric properties were most consistently associated with the predicted haptic-saliency distribution.

### Predicting perceptual choices from exploration

To test whether the information acquired during active exploration could account for subsequent recognition decisions, we compared the presentation of the reference with the two objects encountered during the comparison phase: the target and the competitor. We considered two types of sampled information: global structure, related to where on the object the fingers contacted the surface, and local surface shape, related to the geometric properties encountered at those contact locations. For each type of information, the reference was compared independently with the target and competitor, and the comparison yielding the smaller distance was taken as the model’s predicted response. In addition to model integration of information over the complete exploration, we tested several temporally restricted variants to determine whether recognition might depend disproportionately on information sampled at particular stages of the exploration.

#### Global shape similarity

Contact events were identified from the fingertip trajectories using the same contact criterion described above, and each contact sample was associated with its nearest location on the corresponding pepper surface. To permit comparison across presentations, contacted locations were represented relative to the object rather than in external motion tracker coordinates. In Experiment 1, in which peppers were presented in a fixed orientation, coordinates were centred on the object and normalised by object scale. In Experiment 2, in which object orientation varied across presentations, the known presentation orientation was additionally used to transform the object and its contacted locations into a common canonical orientation before comparison. Distance between two sets of contacted locations was quantified using a symmetric nearest-neighbour measure: the Euclidean distance from each location in the first set to its nearest location in the second set was averaged, the corresponding mean was calculated in the reverse direction, and the two values were averaged. The comparison object producing the smaller symmetric distance to the initial target was selected as the model prediction.

#### Local surface-shape similarity

Each contacted surface location was characterised using the same six curvature-based descriptors used in the geometric analysis of haptic saliency (Figure 2). Local surface geometry was estimated from the 30 nearest surface neighbours.

Neighbourhood coordinates were expressed in a local tangent frame obtained by principal-component analysis, and a quadratic surface was fitted to the local patch to estimate curvature. Because the descriptors differed in numerical scale, feature values were standardised within each trial using the pooled feature distributions over the complete surfaces of the initial target, repeated target and competitor. The resulting six-dimensional feature vectors sampled from the initial target and each comparison object were compared using the same symmetric nearest-neighbour distance in feature space.

#### Temporal sampling conditions

For both types of information, *Full* retained all detected contact samples from the reference, target and competitor and therefore provided the unrestricted baseline. Six temporally restricted alternatives were then evaluated, motivated by findings of Metzger and colleagues (23–25). *Full target → Q1* (first quarter) retained the complete initial-target exploration but only the first 25% of the exploration of the comparison objects. *Full target → half* likewise retained the complete reference exploration but the first 50% of each comparison exploration. *Final → full* retained only the final contact frame from the reference and compared it with the complete comparison explorations. *First only* retained the first contact frame from all three presentations, whereas *Final* only retained the final contact frame from all three. Finally, *Final → first* comparison compared the final contact frame of the initial target with the first contact frame from the comparisons, providing the temporally closest comparison across successive object presentations (highest temporal proximity). For first- and final-frame conditions, all fingertips contacting the surface during the selected frame were retained. For the first-quarter and first-half conditions, time was normalised separately within each presentation from the first detected contact (t = 0) to the final detected contact (t = 1), and all samples within the specified interval were retained. Repeated samples at the 100-Hz tracking rate were preserved. The same symmetric nearest-neighbour distance was used for every temporal condition, so differences between models reflected the information retained rather than a change in the comparison rule.

#### Model performance and statistical analysis

Objective accuracy was the proportion of trials on which the model selected the target and therefore identified the objectively correct object. Choice agreement was the proportion of trials on which the model selected the same comparison object as the participant, irrespective of whether that response was correct. Error agreement was calculated only on trials on which the participant responded incorrectly and was defined as the proportion on which the model also selected the competitor. An error-agreement value below 0.5 therefore indicates that, on human-error trials, the model more often favoured the objectively correct target. All measures were calculated separately for each participant. The temporal analysis was restricted to three theoretically motivated sampling conditions. The Full condition, which used all available contact information from the initial target and both subsequent objects, provided the full-information benchmark. The two restricted conditions were motivated by the temporal pattern reported by Metzger and colleagues (23). In that study, temporal effects were found for information acquired from the subsequently explored objects, whereas no corresponding temporal effect was detected during exploration of the first stimulus. Because the latter finding reflects a null effect rather than positive evidence for temporal invariance, we considered two plausible implementations of this pattern. In the Full → Q1 condition, the complete exploration history of the initially explored target was retained, while the two subsequently explored objects were restricted to the first quartile of their contact interval. This implements the possibility that information from the first stimulus is retained relatively broadly, whereas recognition depends preferentially on early information from later objects. In the Final → first condition, only the final contact state of the initial target was compared with the first contact state of each subsequent object. This provides a stricter temporal-proximity test in which the most recent information from the remembered target is compared with the earliest incoming information from the subsequent objects. Statistical inference was therefore limited to two planned paired-sample comparisons for each dependent measure and information type: Full versus Full → Q1 and Full versus Final → first. Both comparisons tested theoretically motivated temporal restrictions against the same full-information benchmark; no other pairwise temporal comparisons were performed. The two planned tests constituted the correction family for each dependent measure and information type, and significance was assessed using a Bonferroni-corrected threshold of α = .025. Group values are reported as participant means ± SEM.

## Results

### Human accuracy

In Experiment 1, participants identified the matching object with a mean accuracy of 87.1% (SD = 7.6%), significantly above the 50% chance level, t(8) = 14.60, p < .001. In Experiment 2, recognition accuracy was lower but remained significantly above chance (M = 72.7%, SD = 13.6%), t(9) = 5.27, p < .001. Recognition accuracy was significantly higher in Experiment 1 (M = 87.1%, SD = 7.6%) than in Experiment 2 (M = 72.7%, SD = 13.6%), t (17) = 2.80, p = .012.

### Predicting haptic saliency — Experiment 1

The geometry-based model generalised to peppers that had been completely excluded from training. Across the ten leave-one-pepper-out folds, mean model performance was AUC=.765, ± .010 SEM, compared with AUC=.743 ± .010 for the generic spatial prior. The geometry model outperformed the generic prior for all ten held-out peppers (Fig. 3A), with a mean improvement of .0225 AUC units, t(9)=6.33, p<.001. Thus, local surface geometry predicted where participants touched over and above spatial regularities shared across peppers.

**Figure 3.**
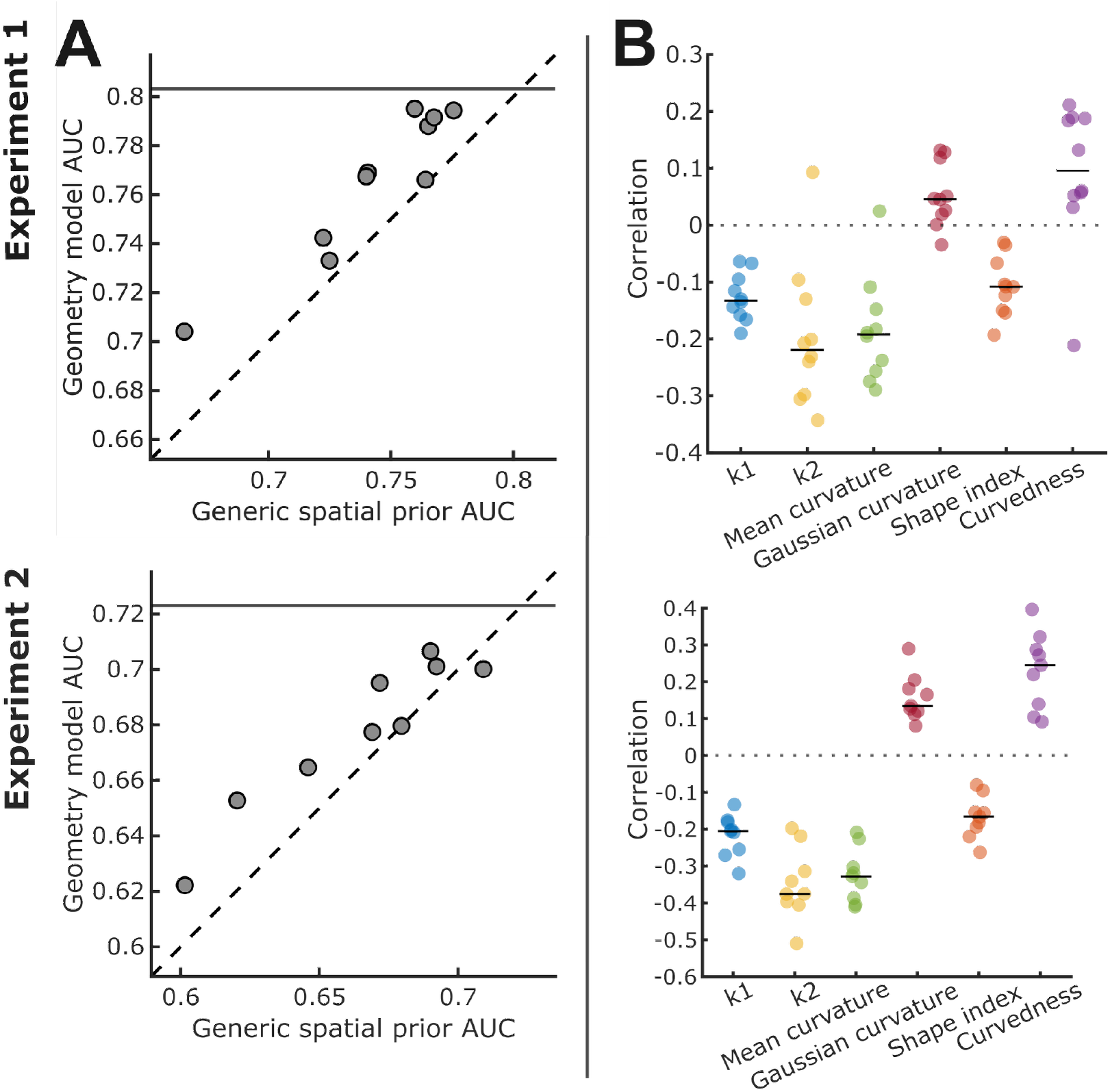
Geometry-based prediction of haptic saliency. Experiment 1 (top), Experiment 2 (bottom) **(A)** Predictive performance for each held-out pepper. Each point shows the AUC of the geometry model plotted against the AUC of the generic spatial prior for the same pepper. The dashed diagonal indicates equal performance of the two models; points above this line indicate better prediction by the geometry model. The solid horizontal line shows the mean same-pepper participant-agreement AUC, providing an empirical upper reference for model performance. **(B)** Relationship between model-predicted haptic saliency and local surface geometry. Each point represents one pepper, after collapsing estimates across the 24 rigid rotations by their median; horizontal black bars indicate the median correlation across peppers. The dashed horizontal line indicates zero correlation.

The same-pepper participant-agreement reference reached AUC=.803 ± .010 SEM. Model performance remained lower than this empirical upper reference by .038 AUC units, *t*(9)=6.50, *p*<.001, indicating that geometry accounted for only part of the reproducible object-specific structure in human exploration.

We next examined which aspects of local surface geometry were associated with the saliency predicted by the network. For each pepper, Spearman correlations between predicted saliency and each geometric descriptor were computed across the surface and then summarised across peppers (Fig. 3B). Predicted saliency was negatively associated with principal curvature k1 (median ρ=-.132, *t*(9) = - 9.62, *p*<.001), principal curvature k2 (median ρ=-.219, *t*(9)=-4.88, *p*<.001), mean curvature (median ρ=-.192, *t*(9)=-6.30, *p*<.001), and shape index (median ρ=-.108, *t*(9)=-6.52, *p*<.001). Gaussian curvature showed a smaller positive association (median ρ=.046, *t*(9)=2.99, *p*=.015). Curvedness was also positive in direction (median ρ=.096), but did not significantly differ from zero, (*t*(9)=2.24, *p*=.051).

### Predicting haptic saliency — Experiment 2

The same analysis was applied to Experiment 2, in which peppers were presented in different orientations. The geometry model again generalised to held-out objects. Across the nine leave-one-pepper-out folds, mean performance was AUC=.678 ± .009 SEM, compared with AUC=.664 ±.012 SEM for the generic spatial prior. The model outperformed the generic prior for eight of the nine peppers (Fig. 4A), with a mean improvement of .0133 AUC units, *t(*8)=3.17, *p*=.013. Participant agreement reached AUC=.723 ± .006, exceeding model performance by .045 AUC units, *t*(8)=5.74, *p*<.001. Thus, even when object orientation varied across trials, local geometry explained reproducible object-specific variation in touch location beyond the generic spatial prior, while remaining below the consistency observed between participants exploring the same object. Predicted saliency was consistently related to local geometry across peppers in Experiment 2 (Fig. 4B), with all six geometric measures differing significantly from zero. Predicted saliency was negatively associated with principal curvature k1 (median ρ =-.205, *t*(8)=-11.53, *p*<.001), principal curvature k2 (median ρ =-.376, *t*(8)=-10.87, *p*<.001), mean curvature (median ρ =-.328, *t*(8)=-13.52, *p*<.001), and shape index (median ρ =-.166, *t*(8)=-8.85, *p*<.001). Gaussian curvature was positively associated with predicted saliency (median ρ =.135, *t*(8)=7.55, *p*<.001), as was curvedness (median ρ =.245, *t*(8)=6.73, *p*<.001). Across the two experiments, the model therefore showed the same qualitative geometric signature: negative associations with the two principal curvatures, mean curvature and shape index, together with a positive association with Gaussian curvature. The positive association with curvedness was clear in Experiment 2 but weaker and not reliable in Experiment 1. This pattern suggests that haptic saliency is related to a particular combination of signed local surface curvatures rather than simply to the overall magnitude of local bending.

**Figure 4.**
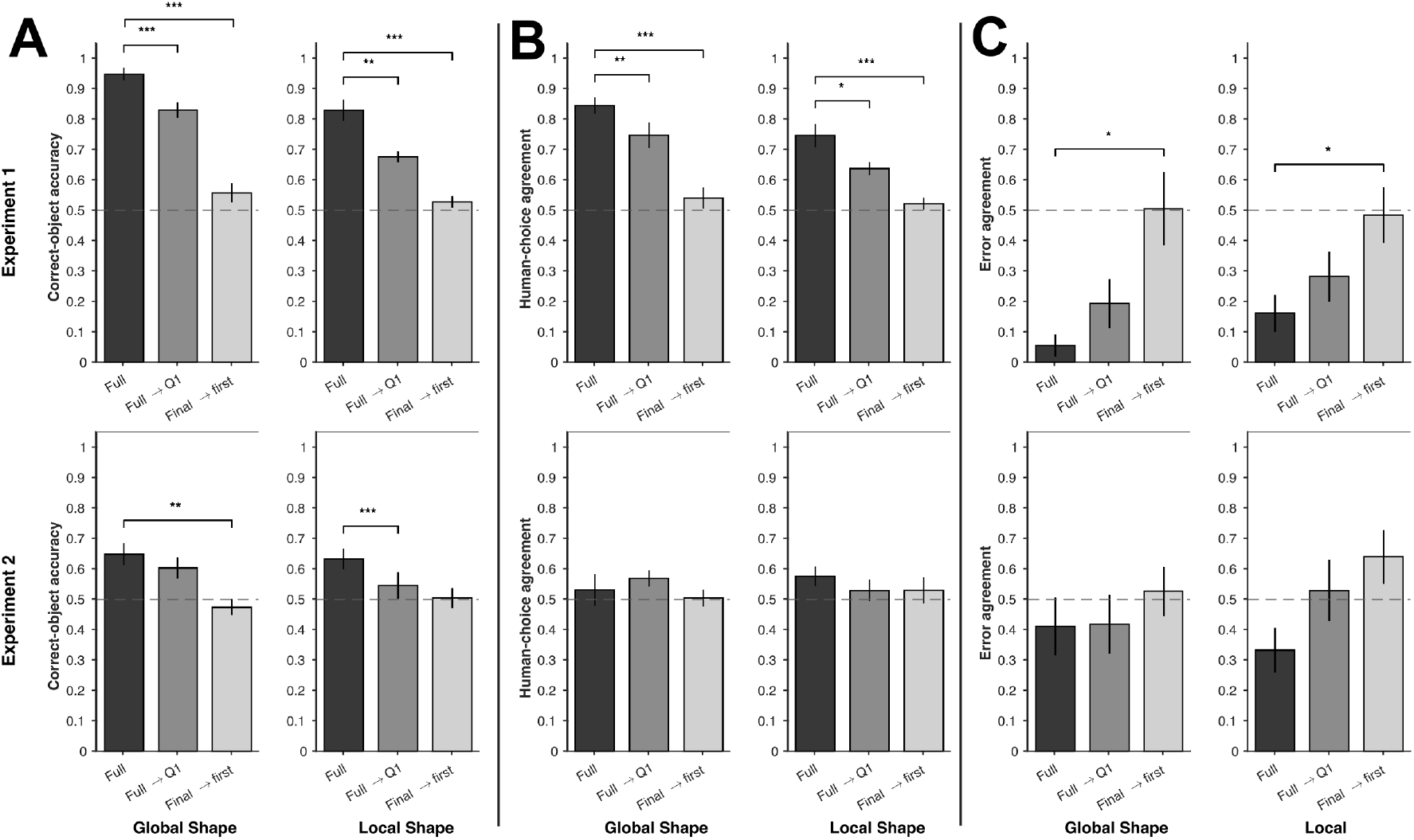
Prediction of perceptual choices. Model performance based on global and local shape information for the three sampling models in Experiment 1 (top) and Experiment 2 (bottom). (A) Correct-object accuracy. (B) Human-choice agreement. (C) Error agreement. Bars show participant means ± SEM; the dashed horizontal line indicates chance performance (0.5). Brackets indicate the two prespecified comparisons with the Full condition that were significant at the Bonferroni-corrected threshold of α = .025. If asterisks are used, * denotes p < .025, ** p < .01, and *** p < .001.

### Predicting recognition choices from exploration, Experiment 1 and 2

In Experiment 1 (Figure 4, top panel), the Full exploration model contained strong information about the objectively correct object, with correct-object accuracy of .948 ± .021 for global shape and .829 ± .035 for local shape. Choice agreement was also high (.845 ± .027 and .747 ± .038, respectively), consistent with the high recognition performance of participants, whereas error agreement was low (.056 ± .037 and .162 ± .061). Thus, when all sampled information was available, the model generally continued to favour the objectively correct pepper even on trials on which participants responded incorrectly. Correct-object accuracy showed significant effects of information type, F(1, 8) = 30.43, p < .001, η_p_ ^2^ = .792, and temporal sampling condition, F(2, 16) = 63.04, p < .001, ηp2 = .887, together with an information type × temporal condition interaction, F(2, 16) = 3.77, p = .045, η _p_ ^2^ = .321. For global shape, accuracy decreased relative to **Full** exploration for both **Full → Q1**, t(8) = −6.77, p < .001, and Final → first, t(8) = −8.66, p < .001. The same pattern was observed for local shape: accuracy decreased for **Full → Q1**, t(8) = −4.46, p = .002, and **Final → first**, t(8) = −6.73, p < .001. All four planned comparisons survived the Bonferroni-corrected threshold of α = .025. Choice agreement likewise showed significant effects of information type, F(1, 8) = 8.58, p = .019, η _p_ ^2^ = .518, and temporal sampling condition, F(2, 16) = 31.80, p < .001, η _p_ ^2^ = .799, with no reliable information type × temporal condition interaction. For global shape, choice agreement decreased relative to **Full** exploration for **Full → Q1**, t(8) = −4.39, p = .002, and **Final → first**, t(8) = −7.31, p < .001. For local shape, choice agreement was also lower for **Full → Q1**, t(8) = −3.26, p = .012, and **Final → first**, t(8) = −5.12, p = .001. Again, all planned comparisons were significant. Error agreement showed a significant effect of temporal sampling condition, F(2, 16) = 8.21, p = .004, η_p_ ^2^ = .506, but no effect of information type, and no interaction. **Full → Q1** did not significantly differ from **Full** exploration for either global shape, t(8) = 1.40, p = .200, or local shape (.162 vs. .282), t(8) = 2.29, p = .052. In contrast, **Final → first** increased error agreement to .506 for global shape, t(8) = 3.15, p = .014, and to .484 for local shape, t(8) = 2.77, p = .024. Thus, the strongest temporal proximity substantially reduced objective identification while simultaneously making the model more consistent with participants’ errors.

Experiment 2 (Figure 4, bottom panel) showed a weaker but convergent pattern. Under **Full** exploration, correct-object accuracy was .649 ± .037 for global shape and .632 ± .034 for local shape. Temporal sampling condition significantly affected correct-object accuracy, F(2, 18) = 10.73, p < .001, η_p_ ^2^ = .544, whereas neither information type nor the interaction were significant. For global shape, Final → first reduced accuracy relative to **Full** exploration, t(9) = −3.54, p = .006, whereas **Full → Q1** did not. For local shape, **Full → Q1** reduced accuracy, t(9) = −5.19, p < .001, whereas the reduction for **Final → first**, t(9) = −2.33, p = .045, did not meet the corrected α = .025 criterion but showed a trend. Choice agreement did not vary reliably with information type temporal sampling condition or their interaction. Error agreement showed a significant overall effect of temporal sampling condition, F(2, 18) = 4.33, p = .029, η_p_^2^= .325=, with no effect of information type and no interaction. For global shape, error agreement did not differ between **Full** exploration and either **Full → Q1**, or **Final → first**. For local shape, error agreement increased descriptively from .333 under **Full** exploration to .529 for **Full → Q1** and .640 for **Final → first**, but neither planned comparison reached the Bonferroni-corrected threshold of α = .025: **Full → Q1**, t(9) = 1.72, p = .120, and **Final → first**, t(9) = 2.27, p = .049.

Taken together, the analyses indicate that temporal sampling systematically altered the information available for recognition. Full exploration generally provided the strongest information about the objectively correct object, whereas restricting the temporal information weakened objective identification. In Experiment 1, the Final → first model additionally produced a reliable increase in agreement with participants’ errors, demonstrating a clear trade-off between objective identification and compatibility with human errors. Experiment 2 showed the same qualitative tendency at the omnibus level: temporal sampling significantly affected both objective accuracy and error agreement, although no individual error-agreement contrast survived correction. The results are therefore consistent with recognition decisions being influenced by temporal weighting of acquired haptic information rather than by an optimal integration of the complete exploration history.

## Discussion

The present study examined how active touch samples the surface of fully three-dimensional objects and whether the information acquired through this sampling can account for subsequent perceptual choices. Three main findings emerged. First, haptic exploration was highly structured. A substantial component of touch behaviour was shared across different peppers, but local object geometry provided additional object-specific information: the geometry-based network predicted where participants touched on previously unseen peppers better than the generic spatial prior in both experiments. Second, predicted saliency showed a consistent relationship with local three-dimensional geometry. Across experiments, touch was associated with a particular combination of signed curvature measures rather than simply with greater overall curvature magnitude. Third, the information acquired during exploration was highly informative about object identity, but participants did not appear to use all sampled information in an equally weighted or optimal manner. Models integrating the complete exploration often identified the objectively correct pepper even when participants responded incorrectly, whereas restricting the comparison to temporally restricted portions of the exploration reduced objective identification but made predictions more compatible with participants’ errors. This pattern suggests that recognition may rely disproportionately on information sampled close in time across successive object presentations, potentially because of memory decay. Thus, the behavioural decision appears to reflect not simply the information acquired during exploration, but how that information is retained and weighted over time.

A striking feature of the touch distributions was the strong pepper-aspecific spatial component. This is consistent with the broader literature showing that haptic exploration is far from arbitrary. Manual exploration typically follows systematic exploratory procedures that are adapted to both the task and the information being sought (2,12). Previous work on haptic saliency likewise found substantial stimulus-independent regularities in where people touch, alongside systematic effects of local stimulus properties (3,32). Our results extend this principle from relief-like surfaces to three-dimensional objects. Although much of the exploration pattern could be predicted without knowing which pepper was being touched, local geometry nevertheless improved prediction for peppers that had been entirely excluded from model training. Haptic saliency therefore appears to combine generic sensorimotor regularities with stimulus-dependent selection.

The generic component should, however, be interpreted partly in light of the stimulus configuration used here. In Experiment 1, the peppers had a fixed orientation, and in Experiment 2 orientation varied only by rotation around the vertical axis. Thus, the relation of the object to gravity, its upper and lower regions, the support surface and many aspects of the observer’s approach remained relatively stable. A stronger test of feature based haptic saliency would rotate objects around multiple axes and vary initial hand posture or object support. Importantly, however, this does not undermine the object-specific effect observed here: in both experiments, local geometry improved prediction beyond the generic spatial prior.

The geometric analysis further indicates that haptic saliency cannot be reduced to a simple preference for highly curved regions.

Predicted saliency was negatively associated with both principal curvature estimates, mean curvature and shape index, and positively associated with Gaussian curvature in both experiments; curvedness was clearly positive only in Experiment 2. Importantly, a complementary analysis of the observed human touch distributions showed a similar qualitative relationship with these curvature measures (Supplementary data), indicating that the geometric signature recovered by the network reflects corresponding structure in participants’ actual sampling behaviour. This further supports a role for local curvature in haptic saliency. The combination of positive Gaussian curvature with negative principal and mean curvatures indicates preferential sampling of locally convex, dome-like surface regions rather than saddle-like regions. By contrast, curvedness, which measures curvature magnitude irrespective of sign, showed a weaker and less consistent relationship with touch. Thus, haptic saliency appears to depend on the type of local shape, rather than simply on how strongly the surface is curved. This complements recent findings from 2D contour exploration, where observers preferentially explored particular geometric features, especially concavities (33). Although contour concavity and 3D surface curvature are not directly equivalent, both findings suggest that active touch selectively prioritises specific geometric configurations rather than curvature magnitude alone The recognition analyses reveal an important dissociation between the information contained in exploration and the information apparently used for the final decision. In Experiment 1, the model based on the complete exploration identified the objectively correct object with very high accuracy, particularly from global shape information, yet their agreement with participants on error trials was extremely low. In other words, the locations and local shape properties that participants themselves had sampled frequently contained sufficient information to identify the target among the comparison objects, even on trials in which they chose the competitor. This argues against a simple account in which behavioural errors arise solely because participants failed to touch informative parts of the object. The temporal analyses support this interpretation. Using restricted portions of the exploration reduced the model’s ability to recover the objectively correct pepper. When only the final contact state of the reference was compared with the first contact state of the comparison objects accuracy is reduced while error agreement increased towards chance for both contact position and local shape. This suggests that behavioural decisions may reflect a temporally weighted subset of the information that was physically acquired.

This temporal pattern is consistent with previous work on softness discrimination (23,34). Participants explored two stimuli sequentially using repeated indentations: estimates obtained from the first stimulus contributed relatively uniformly to the judgement, whereas later estimates from the second stimulus contributed progressively less, probably due to memory constraints. Our findings support the idea that memory limitations influence the temporal integration of haptic information. However, the present analyses do not identify the mechanism responsible for this temporal weighting. Furthermore, it remains unclear if participants are at all capable to construct a coherent representation of the object which would be required for integration throughout a prolonged exploration.

The use of bell peppers also defines the scope of our conclusions. These stimuli are advantageous because exemplars share the same semantic category and volume while differing substantially in idiosyncratic three-dimensional geometry (26–28). Recognition therefore cannot be solved simply by category membership or size comparison, and differences in exploration can be related directly to shape. At the same time, peppers have little variation in semantic meaning or functional affordance. Exploration of tools and manipulable objects is likely to be influenced by the potential functionality of the object and which regions are relevant to action: exploration of such objects becomes strongly directed towards properties known to be diagnostic for classification (35). For functionally meaningful objects, handles, graspable regions or other action-relevant parts might therefore attract touch even when they are not locally salient in purely geometric terms. Whether the present geometric saliency effects generalise to semantically distinct and affordance-rich objects remains an important question.

The recognition models also deliberately simplify the sensory information available in active touch. Contact-position similarity uses where the fingers contacted the surface but not force, pressure, skin deformation or the full configuration of the hand. The local-shape model represents curvature at sampled positions but does not encode how those features were encountered through movement.

Human recognition may depend on configural relations among sampled features, proprioceptive information available from the whole hand, force and movement history, as well as on memory for the sequence itself. Importantly, our findings provide the first insights and the methodological advances toward the understanding of the relationship between sensory input, internal representation and perceptual decisions.

Finally, the two experiments differed in the strength of several effects. Geometry predicted haptic saliency in both experiments, but participants’ accuracy and recognition effects were generally clearer in the fixed-orientation experiment. This may reflect the additional representational demands imposed by changes in viewpoint in Experiment 2. Haptic object recognition is not completely orientation invariant, and changes in presentation orientation can alter the relation between local object structure, hand posture and previously sampled information (36). The weaker recognition effects in Experiment 2 are therefore compatible with the idea that establishing correspondence across changing object orientations adds noise or requires an additional transformation before comparison. Importantly, however, the central qualitative pattern remained: integrating more sampled information supported objective identification, whereas more temporally proximal comparisons were relatively more compatible with the pattern of behavioural errors.

In summary, active haptic exploration of three-dimensional objects is shaped by both generic sensorimotor regularities and object-specific surface geometry. Local geometry predicts where people touch beyond a strong pepper-aspecific exploration bias, and the resulting samples contain considerable information about object identity. Recognition, however, is not a simple read-out of all of this information. Participants can make errors even when their exploration contains sufficient information for a simple model to recover the correct object, and temporally proximal comparisons are more compatible with those errors than full integration of the exploration. Haptic saliency and haptic recognition should therefore be treated as related but distinct stages: object geometry helps determine what information is acquired, whereas temporal weighting, memory and comparison processes help determine which parts of that information ultimately guide the decision.

## Acknowledgements

The research was supported by the Royal Society RG\R1\241159 – Research Grant. We thank J. Farley Norman for sharing the 3D models of the peppers with us.

## References

1. Gibson JJ. Observations on active touch. Psychol Rev. 1962;69(6):477.

2. Lederman SJ, Klatzky RL. Hand movements: A window into haptic object recognition. Cognit Psychol. 1987;19(3):342–68.

3. Itti L, Koch C, Niebur E. A model of saliency-based visual attention for rapid scene analysis. IEEE Trans Pattern Anal Mach Intell. 1998;20(11):1254–9.

4. Hayhoe M, Ballard D. Modeling task control of eye movements. Curr Biol. 2014;24(13):R622–8.

5. Ballard DH, Hayhoe MM. Modelling the role of task in the control of gaze. Vis Cogn. 2009;17(6–7):1185–204.

6. Toscani M, Yücel EI, Doerschner K. Gloss and speed judgments yield different fine tuning of saccadic sampling in dynamic scenes. - Percept. 2019;10(6):2041669519889070.

7. Metzger A, Ennis RJ, Doerschner K, Toscani M. Perceptual task drives later fixations and long latency saccades, while early fixations and short latency saccades are more automatic. Perception. 2024;53(8):501–11.

8. Metzger A, Toscani M, Valsecchi M, Drewing K. Haptic Saliency Model for Rigid Textured Surfaces. In: Prattichizzo D, Shinoda H, Tan HZ, Ruffaldi E, Frisoli A, editors. Haptics: Science, Technology, and Applications. Cham: Springer International Publishing; 2018. p. 389–400. (Lecture Notes in Computer Science). doi:10.1007/978-3-319-93445-7_34

9. Metzger A, Toscani M, Akbarinia A, Valsecchi M, Drewing K. Deep neural network model of haptic saliency. Sci Rep. 2021 Jan 14;11(1):1395. doi:10.1038/s41598-020-80675-6

10. Najemnik J, Geisler WS. Optimal eye movement strategies in visual search. Nature. 2005;434(7031):387–91.

11. Toscani M, Valsecchi M, Gegenfurtner KR. Optimal sampling of visual information for lightness judgments. Proc Natl Acad Sci. 2013;110(27):11163–8.

12. Lederman SJ, Klatzky RL. Extracting object properties through haptic exploration. Acta Psychol (Amst). 1993;84(1):29–40.

13. Lederman SJ, Klatzky RL. Haptic identification of common objects: Effects of constraining the manual exploration process. Percept Psychophys. 2004;66(4):618–28.

14. Metzger A, Toscani M, Valsecchi M, Drewing K. Foveation-like behavior in human haptic search. J Vis. 2020 Oct 20;20(11):1105. doi:10.1167/jov.20.11.1105

15. Berryman LJ, Yau JM, Hsiao SS. Representation of Object Size in the Somatosensory System. J Neurophysiol. 2006 Jul;96(1):27–39. doi:10.1152/jn.01190.2005

16. Hsiao S. Central mechanisms of tactile shape perception. Curr Opin Neurobiol. 2008 Aug;18(4):418–24. doi:10.1016/j.conb.2008.09.001

17. Voisin J, Lamarre Y, Chapman CE. Haptic discrimination of object shape in humans: contribution of cutaneous and proprioceptive inputs. Exp Brain Res. 2002 Jul;145(2):251–60. doi:10.1007/s00221-002-1118-5

18. Yau JM, Kim SS, Thakur PH, Bensmaia SJ. Feeling form: the neural basis of haptic shape perception. J Neurophysiol. 2016 Feb 1;115(2):631–42. doi:10.1152/jn.00598.2015

19. Kappers AM. Human perception of shape from touch. Philos Trans R Soc B Biol Sci. 2011;366(1581):3106.

20. Kappers AM, Koenderink JJ, Pas SF te. Haptic discrimination of doubly curved surfaces. Perception. 1994;23(12):1483–90.

21. Kappers AM, Koenderink JJ, Lichtenegger I. Haptic identification of curved surfaces. Percept Psychophys. 1994;56(1):53–61.

22. Newell FN, Ernst MO, Tjan BS, Bülthoff HH. Viewpoint Dependence in Visual and Haptic Object Recognition. Psychol Sci. 2001 Jan;12(1):37–42. doi:10.1111/1467-9280.00307

23. Metzger A, Lezkan A, Drewing K. Integration of serial sensory information in haptic perception of softness. J Exp Psychol Hum Percept Perform. 2018;44(4):551.

24. Metzger A, Drewing K. The longer the first stimulus is explored in softness discrimination the longer it can be compared to the second one. In. IEEE; 2017. p. 31–6.

25. Metzger A, Drewing K. A Kalman filter model for predicting discrimination performance in free and restricted haptic explorations. In. IEEE; 2021. p. 439–44.

26. Norman JF, Norman HF, Clayton AM, Lianekhammy J, Zielke G. The visual and haptic perception of natural object shape. Percept Psychophys. 2004;66(2):342–51.

27. Norman JF, Phillips F, Holmin JS, Norman HF, Beers AM, Boswell AM, et al. Solid shape discrimination from vision and haptics: Natural objects (Capsicum annuum) and Gibson’s “feelies”. Exp Brain Res. 2012;222(3):321–32.

28. Norman JF, Wheeler SP, Pedersen LE, Shain LM, Kinnard JD, Lenoir J. The recognition of solid object shape: the importance of inhomogeneity. -Percept. 2019;10(4):2041669519870553.

29. Hartmann F, Maiello G, Rothkopf CA, Fleming RW. Estimation of Contact Regions Between Hands and Objects During Human Multi-Digit Grasping. J Vis Exp. 2023 Apr 21;(194):64877. doi:10.3791/64877

30. Qi CR, Su H, Mo K, Guibas LJ. Pointnet: Deep learning on point sets for 3d classification and segmentation. In. 2017. p. 652–60.

31. Qi CR, Yi L, Su H, Guibas L. Pointnet++: Deep hierarchical feature learning on point sets in a metric space. Adv Neural Inf Process Syst. 2017;30.

32. Metzger A, Toscani M, Valsecchi M, Drewing K. Haptic saliency model for rigid textured surfaces. In. Springer; 2018. p. 389–400.

33. Lin LP, Kshirsagar A, Belousov B, Schneider T, Peters J, Doerschner K, et al. Exploration Strategies and Feature Prioritisation in Contour-based Haptic Perception of 2D Shape. IEEE Trans Haptics. 2026.

34. Metzger A, Drewing K. Effects of stimulus exploration length and time on the integration of information in haptic softness discrimination. IEEE Trans Haptics. 2019;12(4):451–60.

35. Lederman SJ, Klatzky RL. Haptic classification of common objects: Knowledge-driven exploration. Cognit Psychol. 1990;22(4):421–59.

36. Newell FN, Ernst MO, Tjan BS, Bülthoff HH. Viewpoint dependence in visual and haptic object recognition. Psychol Sci. 2001;12(1):37–42.

