## Supplementary data for "From haptic exploration to object perception in 3D"

### Supplementary analysis: relationship between human touch density and local surface geometry

We analysis tested whether the local geometric properties used to interpret the saliency model were also systematically related to where participants actually touched the objects. The analysis was performed separately for Experiments 1 and 2 using the participant touch-density maps and the same six local surface descriptors used in the model analysis.

For each object (pepper), participant-level continuous touch-density maps were first averaged so that each available participant contributed equally to the object-level human touch map. The resulting map therefore described the relative spatial distribution of human haptic sampling over the object surface rather than the absolute number of recorded samples.

At every sampled surface location, six local shape descriptors were computed: the two principal curvatures (k1 and k2), mean curvature, Gaussian curvature, shape index, and curvedness. For each object and descriptor, the association between local human touch density and the local shape value was quantified using Spearman rank correlation across surface points.

The inferential unit was the object. The median correlation for each object was tested against zero across objects using a two-sided one-sample t-test. Experiment 1 contained 10 objects and Experiment 2 contained 9 objects in this analysis. The plotted points in Supplementary Figure X correspond to these object-level median correlations.

### Results

The same qualitative geometric pattern was evident in both experiments. Human touch density was negatively related to principal curvature k1, principal curvature k2, mean curvature, and shape index, whereas Gaussian curvature was positively related to touch density. These associations were significant in both experiments. Curvedness was the exception: it was not reliably associated with touch density in Experiment 1, but showed a weaker positive association in Experiment 2.

#### Supplementary descriptive statistics

| **Shape descriptor** | **Exp. 1 mean rho** | **t(9)** | **p** | **Exp. 2 mean rho** | **t(8)** | **p** |
| --- | --- | --- | --- | --- | --- | --- |
| Principal curvature k1 | -0.302 | -25.60 | < .001 | -0.269 | -11.50 | < .001 |
| Principal curvature k2 | -0.300 | -8.92 | < .001 | -0.299 | -7.24 | < .001 |
| Mean curvature | -0.359 | -16.19 | < .001 | -0.337 | -10.70 | < .001 |
| Gaussian curvature | 0.173 | 13.69 | < .001 | 0.201 | 11.57 | < .001 |
| Shape index | -0.267 | -13.59 | < .001 | -0.228 | -9.55 | < .001 |
| Curvedness | 0.024 | 0.60 | = .561 | 0.129 | 2.48 | = .038 |

These results provide a behavioural validation of the model-based geometric analysis: the spatial distribution of human touch was itself systematically related to local surface geometry. The replication of the same directional pattern across the two experiments indicates that these geometry-touch relationships were not specific to the fixed-orientation condition of Experiment 1.

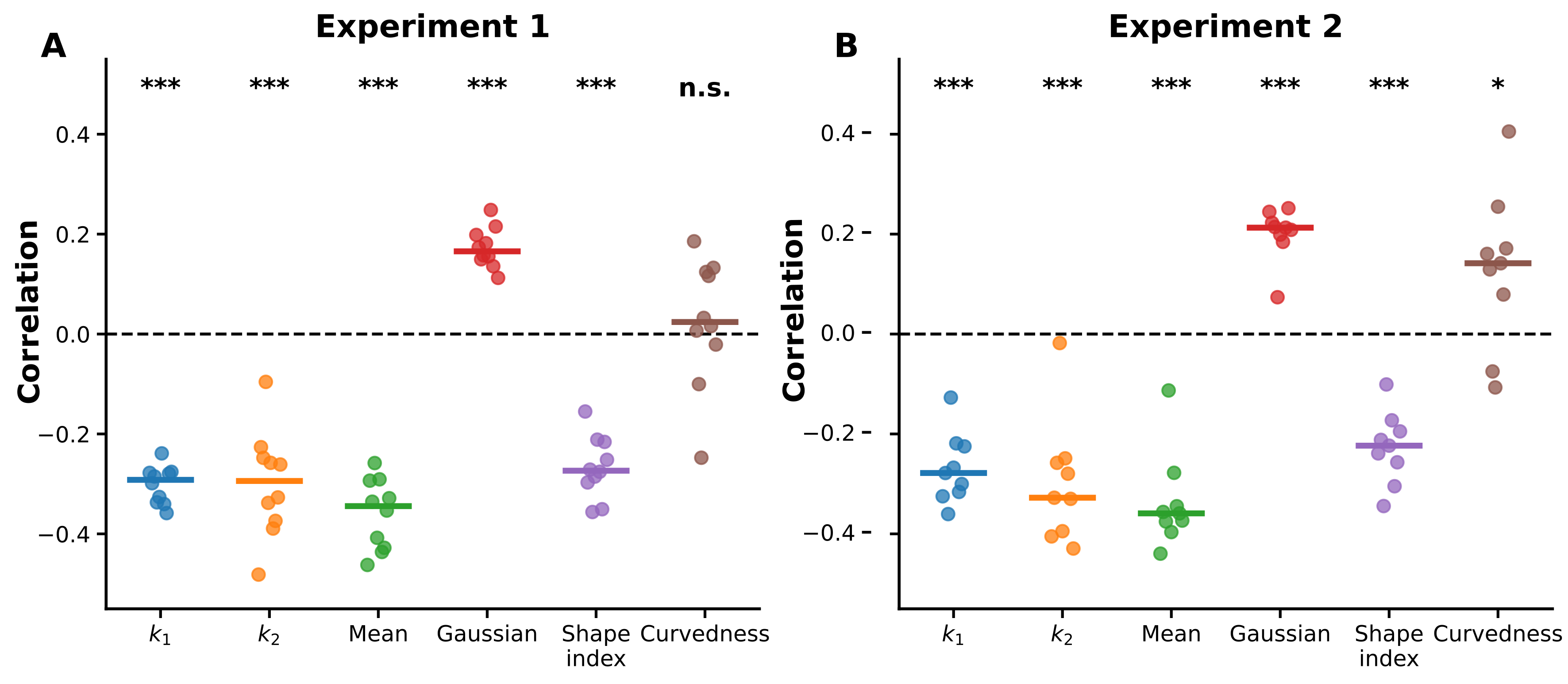

***Supplementary Figure 1. Relationship between local surface geometry and human touch density.*** *Spearman correlations between mean human touch density and six local shape descriptors are shown separately for Experiment 1 (A) and Experiment 2 (B). Each point represents one. Horizontal lines indicate the median across objects, and the dashed horizontal line indicates zero correlation. Both experiments showed the same qualitative pattern: touch density was negatively associated with principal curvature k1, principal curvature k2, mean curvature, and shape index, and positively associated with Gaussian curvature. Curvedness showed no reliable association in Experiment 1 and a weaker positive association in Experiment 2. Significance markers refer to two-sided one-sample t-tests across objects: * p < .05, ** p < .01, *** p < .001.*
